# Amphipathic helices of cellular proteins can replace the helix in M2 of Influenza A virus with only small effects on virus replication

**DOI:** 10.1101/776864

**Authors:** Bodan Hu, Stefanie Siche, Lars Möller, Michael Veit

## Abstract

M2 of influenza virus functions as proton channel during virus entry. In addition, an amphipathic helix in its cytoplasmic tail plays a role during budding. It targets M2 to the assembly site where it inserts into the inner membrane leaflet to induce curvature that causes virus scission. Since vesicularisation of membranes can be performed by a variety of amphiphilic peptides we used reverse genetics to investigate whether they can substitute for M2’s helix.

Virus could not be generated if M2’s helix was deleted or replaced by a peptide predicted not to form an amphiphilic helix. In contrast, viruses could be rescued if the M2 helix was exchanged by helices known to induce membrane curvature. Infectious virus titers were marginally reduced if M2 contains the helix of the amphipathic lipid packing sensor, from the Epsin N-Terminal Homology domain or the non-natural membrane inducer RW16. Transmission EM of infected cells did not reveal unequivocal evidence that virus budding or membrane scission was disturbed in any of the mutants. Instead, individual virus mutants exhibit other defects in M2, such as reduced surface expression, incorporation into virus particles and ion channel activity. The protein composition and specific infectivity was also altered for mutant virions. We conclude that the presence of an amphiphilic helix in M2 is essential for virus replication, but other helices can replace its basic (curvature-inducing) function.

**Importance:** Influenza is unique among enveloped viruses since it does not rely on the cellular ESCRT-machinery for budding. Instead viruses encode their own scission machine, the M2 protein. M2 is targeted to the edge of the viral assembly site where it inserts an amphiphilic helix into the membrane to induce curvature. Cellular proteins utilize a similar mechanism for scission of vesicles. We show that the helix of M2 can be replaced by helices from cellular proteins with only small effects on virus replication. No evidence was obtained that budding is disturbed, but individual mutants exhibit other defects in M2 which explain the reduced virus titers. In contrast, no virus could be generated if the helix of M2 is deleted or replaced by irrelevant sequences. These experiments support the concept that M2 requires an amphiphilic helix to induce membrane curvature, but its biophysical properties are more important than the amino acid sequence.

## Introduction

Influenza A viruses are pleomorphic enveloped viruses in the family *Orthomyxoviridae*. Their membrane is lined from beneath by a protein layer composed of the matrix protein M1, which in turn envelopes the viral genome, arranged as eight viral ribonucleoprotein particles (vRNPs), each composed of a segment of viral RNA complexed to the nucleoprotein (NP) and the subunits of the viral RNA polymerase. The membrane contains three transmembrane proteins, the glycoproteins hemagglutinin (HA), which catalyzes virus entry by receptor-binding and membrane fusion after virus-uptake into endosomes and the neuraminidase (NA), which is required for release of particles from infected cells (1). The tetrameric proton channel protein M2 is activated by the acidic pH in the endosome and the resulting proton flux into the interior of the virus is important for genome unpacking (2–4).

M2 is a type III transmembrane protein; the first 24 amino acids form the ectodomain, which contains an unused glycosylation site and two cysteines which build intermolecular disulphide-linkages, which however, are not required for tetramer formation (5, 6). The following 19 residues are the transmembrane region (TMR), which, including a few amino acids on both sides, is the functional core of the proton channel (7). The remaining 54 residues build up the cytoplasmic tail, which is essential for virus replication (8). Amino acids 48 – 58 following the TMR shape a membrane-parallel amphiphilic helix (9). This region (residues 45-69), but also residues 71–73 contain a binding site for the viral matrix protein M1 (10, 11). The interaction with M2 is required for transport of M1 from internal membranes, where it accumulates in the absence of other virus proteins, to the viral budozone (12, 13), a cholesterol- and sphingomyelin-enriched area of the plasma membrane. The cytoplasmic tail (residues 91 to 94) also binds to the autophagy protein LC3 and recruits LC3-containing vesicles to the plasma membrane, a process that is required for budding of filamentous particles and enhances the environmental stability of virions (14).

In the infected cell, all viral components are synthesized and ultimately transported to the plasma membrane for assembly and budding of progeny virions (15, 16), which occurs in cholesterol- and sphingolipid-enriched nanodomains of the plasma membrane (rafts) (17). HA and NA have intrinsic features that target the proteins to the cholesterol-enriched domains (18), but M2 is not a raft component (19). Nevertheless, M2 co-cluster at the plasma membrane with HA as assessed by FRET and quantitative electron microscopy (20, 21).

It was originally proposed that targeting to the HA-defined budozone is achieved by two features: attachment of palmitic acid to cysteine residue 50 (22, 23), and cholesterol binding (24), mediated by cholesterol recognition/interaction amino acid consensus (CRAC) motifs, present up to four times in the amphiphilic helix region of M2 (25). However, it was then shown that mutations in either the acylation site or in the CRAC motifs, or in both motifs simultaneously do not affect clustering of M2 with HA (26) and virus growth in cell culture (27–29). However, a recent NMR analysis indicated that cholesterol binds the C-terminal part of the transmembrane region and is oriented parallel to the bilayer normal, without requirement of the CRAC motif (30).

Targeting to the budding zone might be achieved by the ability of M2’s helix to sense membrane curvature as shown by preferential binding of M2 to small unilamellar vesicles (SUVs) having a small diameter and by molecular dynamics simulations (31, 32). More specifically, clusters of M2 molecules are excluded from regions with negative curvature, i. e. the outward part of a budding virus particle, but rather accumulate at membrane regions with positive curvature, i. e. the neck of a budding virus (33).

Positioning of M2 at the edge of the viral budding zone could then entail the scission of virus particles: the amphiphilic helix was proposed to induce curvature by wedge-like integration into the membrane (34). Indeed, there is experimental evidence from membrane model system that M2 plays an active role in scission. M2 as well as a peptide encompassing the amphiphilic helix induce membrane curvature in large unilamellar vesicles (LUVs) and causes budding of vesicles into giant unilamellar vesicles (GUVs) (35, 36). Detailed studies with peptides showed that the amphipathic α-helix folds upon contact with membranes. Membrane binding requires hydrophobic interactions with the lipid tails but not charged interactions with the lipid headgroups (31, 32).

In the context of a virus infection it was shown that replacement of residues in the membrane-inserted hydrophobic face of the amphiphilic helix leads to modest or large attenuation of recombinant viruses in cell culture, and to changes in morphology of both filamentous and spherical virus strains. Especially undetached mutant viruses with a bead-on-a-string morphology where individual virions fail to be separated from each other and/or from the plasma membrane are characteristic of incomplete virus scission (36–38).

A multitude of amphipathic helices that interact with membranes have been characterized in cellular proteins. Amphipathic helices are arranged parallel to the bilayer, partially penetrate it via their hydrophobic face and are well suited for membrane deformation and curvature sensing. Their interactions are mostly reversible and restricted to certain cellular membranes in order to allow the respective protein to fulfil a specific cellular function (39). One type of helix induces membrane curvature by inserting into one leaflet of a bilayer like a wedge. A well investigated example is the α0-helix of the Epsin N-Terminal Homology (ENTH) domain, which is present in the Epsin family of proteins that are involved in clathrin-mediated endocytosis. The whole ENTH-domain is composed of several helices that bind specifically to PtdIns(4, 5)P2, a lipid present in the inner leaflet of the plasma membrane. Upon binding to PtdIns(4, 5)P2, the unstructured N-terminal residues of the ENTH domain fold into the α0-helix, that subsequently inserts into the inner leaflet of the plasma membrane resulting in separation of lipid polar heads. The specificity for PtdIns(4, 5)P2 and subsequent membrane insertion is also observed in the isolated α0-helix (40–44).

Instead of inducing membrane curvature, another type of amphipathic helix senses curvature by insertion of bulky hydrophobic residues between loosely packed lipids. The most notable example is the ALPS (amphipathic lipid packing sensor) motif found within the Golgi-associated ArfGAP1 protein. The ALPS motif forms an α-helix only upon binding to highly curved membranes, e.g. to vesicles which have a small radius compared to the rather flat donor membrane. This helix differs from classical amphipathic helices by the abundance of serine and threonine residues on its polar face. Upon lipid-binding ArfGap1 activates the GTPase activity of Arf1, which in turn leads to disassembly of the coat of COPI vesicles, carriers for retrograde transport from the Golgi apparatus to the ER (45–47).

Amphipathic helices can also function as cell-penetrating peptides (CPPs), which are used to deliver membrane impermeable agents through both leaflets of a membrane into cells. The first identified CPPs were composed of only basic amino acids, but peptides containing both basic and hydrophobic amino acids are more effective. Amphipathic CPPs, such as the peptide RW16, bind parallel to the membrane at low concentrations, but at high concentrations insert perpendicularly causing membrane curvature and leakage as well as lipid domain separation and changes in membrane fluidity and cholesterol distribution (48, 49).

Since the mentioned peptides exhibit similar biophysical features and membrane sensing and modulating activities but have a completely different amino acid sequence, we asked whether they can replace the amphiphilic helix of M2 within the context of a virus infection.

## Material and Methods

### Cell culture

Madin Darby canine kidney (MDCK II) cells and human embryonic kidney 293T cells were grown in DMEM (Dulbecco’s Modified Eagle Medium, PAN, Aidenbach, Germany) supplemented with 10% FCS (fetal calf serum, Perbio, Bonn, Germany), 100 units/ml penicillin and 100 μg/ml streptomycin at 37 °C and 5% CO_2_.

### Mutagenesis, generation of recombinant virus, growth curves

Recombinant influenza A/WSN/33 (H1N1) virus was generated using an eight-plasmid reverse genetics system (50), where each plasmid contains the cDNA of one viral RNA segment, flanked by suitable promoters. In brief, 293T cells in 60mm dishes were co-transfected with 8 plasmids (1µg each) with TurboFect transfection reagent. 4-6h post transfection, culture medium was changed to infection medium (DMEM) supplemented with 0.1% FBS, 0.2% BSA and 1 µg/ml TPCK-Trypsin). 48h post transfection, culture supernatant was harvested, centrifuged at 4000g for 5min to clear from cell debris and stored at −80°C or directly passaged onto MDCK II cells for further amplification.

To generate mutations in the amphiphilic helix of M2 the M1/M2-encoding cDNA segment 7 was digested with StuI and Nae I, and a synthetic cDNA sequence was inserted which does not encode the amino acids 48-62 (FKCIYRRFKYGLKRG) that encompass the amphiphilic helix. The synthetic cDNA contains sites for the restriction enzymes ClaI and BspEI at the beginning and end of the amphiphilic helix, which were used to insert synthetic DNA sequences that encode the internal helix from ArfGAP1 (FLNSAMSSLYSGWSSFTTGASKFAS, M2 ALPS), the curvature sensing α0-helix of the ENTH domain from Epsin 1 (SSLRRQMKNIVHN, M2 Epsin) or the artificial cell-penetrating RW16 peptide (RRWRRWWRRWWRRWRR, M2 RW16). Two of the mutants contained scrambled M2 sequences; one from the WSN virus (KYGCFRYFIKRGKLR, M2 sWSN), the other from the Udorn strain (FFKLGYLEFKIFRGCRH, M2 sUdorn).

To investigate virus growth MDCK II cells were seeded into 6-well or 24-well plates one day before infection so they could be nearly 100% confluent the next day. Cells were then infected with WSN WT or M2 mutants with an m.o.i of 0.001 (for multiple replication cycle) or 1 (for single replication cycle). After binding for 1h, cells were washed once with DMEM and fresh infection medium was added. An aliquot of the cell culture supernatant was harvested at 9h, 23h, 34h and 47h (multiple replication cycle) or at 6h and 9h (single replication cycle) post infection, cleared from cell debris (2000g, 5min) and titrated by plaque-assay.

For the virus samples collected at 34h post infection, the copy number of vRNA segment M and NA was also determined by RT-qPCR. Viral RNA was extracted from the same volume of cell culture supernatant with RTP DNA/RNA Virus Mini Kit (Stratec) according to the instruction. The amount of M and NA segment was determined with one-step RT-qPCR using SensiFAST™ Probe Lo-ROX One-Step Kit (Bioline). The sequences of primers and probe for M segment detection are AGA TGA GYC TTC TAA CCG AGG TCG (forward primer), TGC AAA NAC ATC YTC AAG TCT CTG (reverse primer) and TCA GGC CCC CTC AAA GCC GA (probe). The sequences of primer and probe for NA segment detection are TGG GTC AAT CTG TAT GGT AGT C (forward primer), GCT GCC TTG GTT GCA TAT T (reverse primer) and TGG ATT AGC CAT TCA ATT CAA ACC GGA (probe). The plasmids for reverse genetics were used as standard to calculate the copy number.

To assess stability of virus particles WSN WT and mutant viruses were diluted with infection medium to 200000 PFU/ml. 500 µl virus (i.e. 100.000 PFU) was put into 24-well plates, which were incubated at 37°C and 5% C0_2_. One aliquot was removed every day and titers were determined by plaque assay.

Plaque assay was performed on MDCK II cells in six well plates. Cells were infected with serial 10fold dilutions of virus, incubated for 1h at 37 °C, washed with PBS and overlaid with 2X MEM (Minimum Essential Medium), 1.25% Avicel (FMC BioPolymer), 1% NaHCO3, 0.1% FCS, 0.2% BSA (dissolved in H2O) and 2µg/ml TPCK-trypsin. After incubation for 48h at 37 °C the cell cultures were fixed with 4% PFA, cells were stained with 0.1% crystal violet, and the plaques were counted.

### Virus purification, SDS-PAGE and Western blot

MDCK II cells were infected with WSN WT or mutants at an m.o.i of 0.001. 48h post infection, culture supernatant was harvested, centrifuged at 4000g for 5min to clear from cell debris and viruses were pelleted (100.000g, 2h). Pelleted viruses were then loaded onto a continuous 20-60% sucrose gradient and centrifuged at 100.000g for 4h. Visible virus bands at a density of 35-50% were collected and resuspended in 1X TNE buffer (10 mM Tris, 100 mM NaCl und 1 mM EDTA, pH 7.4) and again pelleted.

For analysis of the viral protein composition, purified viruses were subjected to 12% SDS-PAGE under non-reducing condition and Coomassie staining.

To determine the amount of M2 in virus particles, purified virus preparations were subjected to SDS-PAGE under reducing condition. Gels were blotted onto polyvinylidene difluoride (PVDF) membrane (GE Healthcare). After blocking of membranes (blocking solution: 5% skim milk powder in PBS with 0.1% Tween-20 (PBST)) for 1h at room temperature, they were incubated with either anti-M2 mAb (14C2; Santa Cruz) or anti-M1 mAb (Abcam) overnight at 4°C. After washing (3×10 min with PBST), horseradish peroxidase-coupled secondary antibody (anti-mice or anti-goat, Sigma-Aldrich, Taufkirchen, Germany, 1:5000) was applied for 1h at room temperature. After washing, signals were detected by chemiluminescence using the ECLplus reagent (Pierce/Thermo, Bonn, Germany) and a Fusion SL camera system (Peqlab, Erlangen, Germany). The density of bands was analyzed with Image J software.

### Immunofluorescence microscopy and flow cytometry

To assess the subcellular localization of M2 by immunofluorescence microscopy, MDCK II cells grown in 6-well plates were infected with WSN WT and mutants with a m.o.i. of 1. The cells were fixed 4.5h after infection with 4% (w/v) paraformaldehyde in PBS for 20min, permeabilized with 0.1% Triton X-100 in PBS for 10 min, blocked (3% BSA in PBS for 1h) and incubated with primary anti-M2 mAb14C2 (1:300 in PBS supplemented with 3% BSA, Santa Cruz) for 1h and then with anti-mouse secondary antibody coupled to Alexa Fluor 488 (1:1000, Sigma) for 30min. Washing with PBS (three times, each for 2min) was performed between each step. Pictures were recorded with a Zeiss Axio Vert A1 inverse epifluorescence microscope.

To quantify total and surface expression levels of M2 fluorescence intensities were determined by flow cytometry. MDCK II cells were infected with WSN WT and mutants with a m.o.i of 1, treated with trypsin/EDTA for detaching from the dishes at 4.5h post infection, pelleted (1200g, 5min), resuspended in growth medium for 30min, washed twice with PBS and fixed with 4% formaldehyde for 20min. For analysis of surface M2, cells were directly blocked with 3% BSA in PBS for 30min. For the total expression levels of M2, cells were additionally permeabilized with 0.1% triton X-100 for 10 min before blocking with BSA. Cells were incubated with primary anti-M2 mAb14C2 (1:300 in PBS supplemented with 3% BSA, Santa Cruz) for 1h, and then with anti-mouse secondary antibody coupled to Alexa Fluor 488 (1:1000, Sigma) for 30 min. The total fluorescence intensity was determined for single cells by flow cytometry; at least 100.000 cells were analyzed. Cells with fluorescence intensities below the value determined for mock infected cells were not taken into consideration. From these data the mean fluorescence intensity was calculated and results for total and surface expression were normalized to surface expression of M2 WT in each of three infection experiments.

### M2 proton channel activity assay

293T cells in 6-well plates at ∼80% confluency were transfected with 3µg plasmid encoding eYFP using TurboFect transfection reagent (Thermo Fisher). 24h post transfection, cells were infected with WSN WT or mutants at a moi of 0.5. 16h after infection, cells were detached from plates and divided into two parts. One part was directly measured by flow cytometry to analyze the proton channel activity of M2; the other part was fixed and stained with anti-M2 mAb 14C2 to determine the M2 amount in the plasma membrane of infected cells.

To analyze the proton channel activity, cells were washed twice with DPBS++ (DPBS with calcium and magnesium) and resuspended in DPBS++ at either pH7.2 or pH5.5. The mean fluorescence intensity (MFI) of eYFP of at least 10000 cells was measured every 40s. The MFI at pH7.2 at the starting time point (T=0) was normalized to 100% and its change was plotted against time.

To determine the M2 amount at the plasma membrane, cells were fixed with 4% formaldehyde for 20min, blocked with 3% BSA in PBS for 30min, incubated with primary anti-M2 mAb (1:300 in PBS supplemented with 3% BSA; 14C2, Santa Cruz) for 1h, then with anti-mice secondary antibody Alexa Fluor 488 (1:1000, Sigma) for 30min and analyzed by flow cytometry. The mean fluorescence intensity was calculated from at least 10 000 cells.

### Transmission electron microscopy

MDCK II cells were infected with wild-type and mutant viruses at a m.o.i of 0.001. 24-36h post infection, cells were harvested by scraping, pelleted (2000g, 5min, 4°C), washed twice with HEPES (pH 7.2) and fixed in 2.5% glutaraldehyde in 50 mM HEPES (pH 7.2). After washing in 50 mM HEPES (pH 7.2), cell pellets were embedded in low-melting-point agarose (3% in ddH_2_O, at a ratio of 1:1). Cells were post-fixed with osmium tetroxide (1% in ddH_2_O for 1h), tannic acid [0.1 % in 50 mM HEPES (pH 7.2) for 30min] and uranyl acetate (2 % in ddH_2_O for 2h), dehydrated stepwise in a graded ethanol series and embedded in epon resin. Ultrathin sections (∼60 nm) were prepared with an ultramicrotome (Leica Ultracut UCT) and counterstained with uranyl acetate (2% in ddH_2_O for 20min), followed by lead citrate (Reynolds’ solution for 3min). Ultrathin sections were examined using a JEM-2100 transmission electron microscope (JEOL) at 200 kV. Images were recorded using a Veleta CCD camera (EMSIS).

## Results

### The presence of an amphipathic helix in M2 is essential for virus replication

In order to investigate the role of the amphipathic helix of M2 for replication of the spherical WSN Influenza virus strain (A/WSN 1933, H1N1), residues 48-62 (amino acid sequence FKCIYRRFKYGLKRG) were deleted from the M2 protein.. We also generated four M2 mutants where these amino acid sequences were replaced (Fig. 1). One mutant contained scrambled version of the amphipathic helix of M2 from the WSN virus (KYGCFRYFIKRGKLR, termed M2 sWSN) (36). We also exchanged M2’s helix with three different helices, with the curvature sensing ALPS helix from ArfGAP1 (FLNSAMSSLYSGWSSFTTGASKFAS, M2 Alps), with the curvature inducing α0-helix of the ENTH domain from Epsin 1 (SSLRRQMKNIVHN, M2 Epsin) and with the non-natural cell-penetrating RW16 peptide (RRWRRWWRRWWRRWRR, M2 RW), that was shown to cause budding of vesicles into GUVs, similar to the helix of M2 (36).

**Fig. 1:**
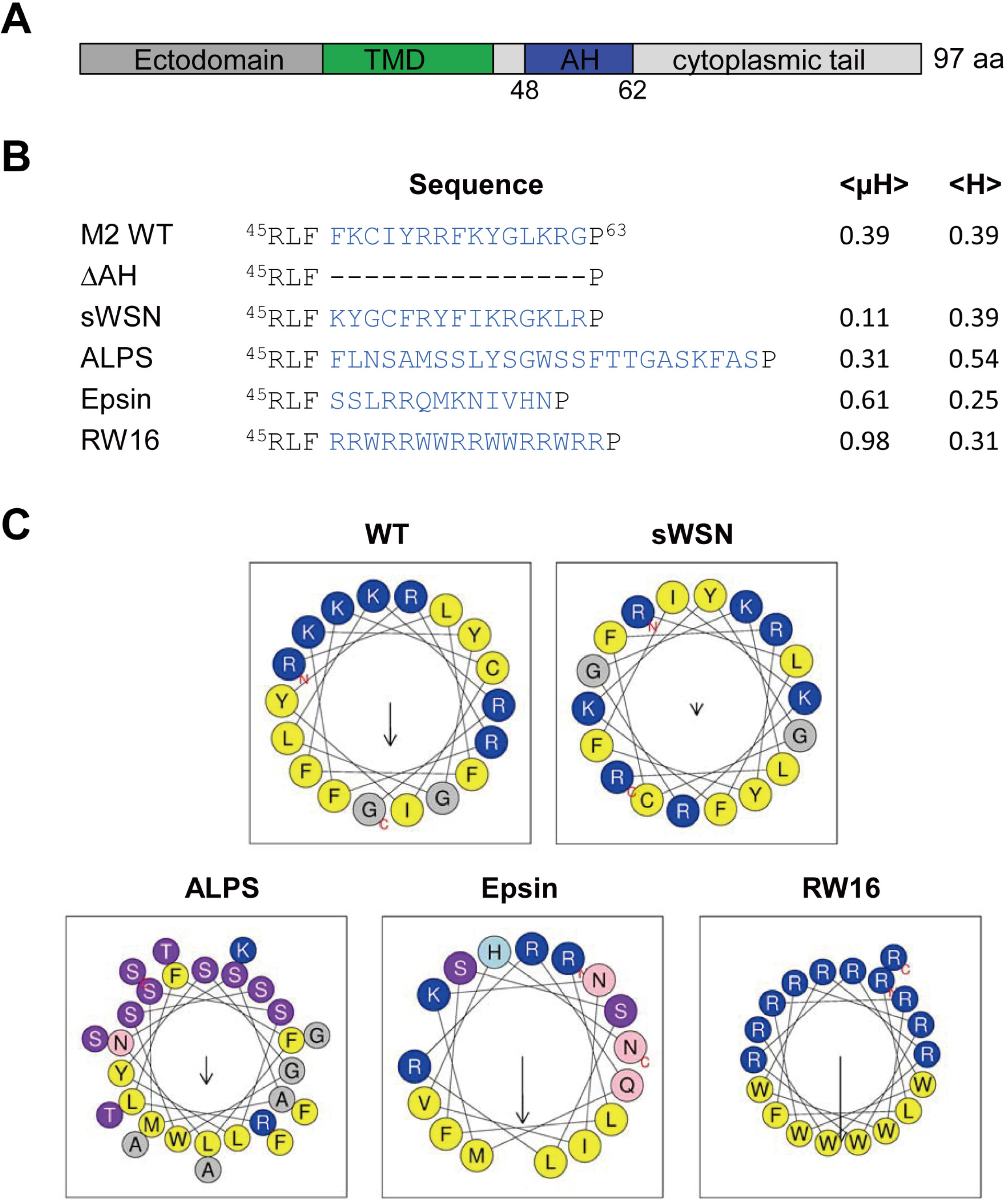
The structure of M2 protein and helical wheel plot of amphipathic helix. (A) Scheme of the M2 protein indicating the individual domains. (B) The amino acid sequences of the amphiphilic region of M2 from the WSN strain and of the mutants investigated in this study. <µH> (hydrophobic moment) and <H> (hydrophobicity) were calculated with heliquest (http://heliquest.ipmc.cnrs.fr/). The scrambled version of the WSN helix was generated with a tool (https://peptidenexus.com/article/sequence-scrambler). (C) Helical wheel plots of the amphiphilic region of M2 WT and the M2 mutants. The arrow points to the hydrophobic face and its length corresponds to the hydrophobic moment. Amino acid sequences used to generate the wheel and the biophysical values in (B) started at R45, but excluded P63.

Amphipathic helices are characterized by two physicochemical parameters, the hydrophobic moment (<µH>) and the average hydrophobicity (<H>). The hydrophobic moment quantifies amphipathicity as the mean vector sum of the hydrophobicities of the side chains if this region forms an α-helix, whereas the hydrophobicity describes the avidity of the helix for lipids (51). Calculation of these parameters by heliquest (Fig. 1c) reveals that M2 WT and M2 sWSN have (since they are composed of identical amino acids) the same hydrophobicity (0.39), but the hydrophobic moment in the scrambled version is reduced from 0.39 to 0.11. M2 ALPS has a similar hydrophobic moment (0.311) as M2 WT and a higher overall hydrophobicity (0.544). M2 Epsin has a higher hydrophobic moment (0.608) compared to M2 WT, but a lower hydrophobicity (0.253). M2 RW16 has the highest hydrophobic moment (0.985) since the peptide is composed of only basic and hydrophobic residues which are perfectly aligned on the hydrophilic and hydrophobic face, respectively.

The mutant M2 plasmid together with plasmids encoding the other viral proteins were transfected into HEK 293T cells, the supernatant was used to infect MDCK II cells and release of virus particles was assessed by HA assay. In five independent transfections we never rescued virus particles which have a deleted helix or the scramble helix of the WSN strain, whereas wild-type virus and the other three mutants done in parallel exhibit HA titers of 2^2^ −2^6^. From the rescued mutants a virus stock was generated in MDCK II cells and sequencing of the M2 gene showed that only the desired mutations were present (data not shown).

To compare the replication kinetics of the viruses, MDCK II cells were infected with WSN WT or with the mutants at a moi of 0.001, supernatants were collected at various time points post infection and virus titers were assessed by plaque assay (Fig. 2A). The growth curve from three infection experiments revealed a small, but statistically significant decrease in the infectious titer of all mutants (depending on the time point) by 1 to 2 logs. To assess whether mutant virus particles exhibit a reduced specific infectivity we also determined their HA-titers at 34 and 47 hours after infection from this experiment (Fig. 2B) and calculated the Pfu/HA ratios at 34 and 47 hours after infection. In general, they are higher at 34 hours compared to 47 hours after infection, which suggests that virus assembly is more precise at earlier time points after infection. More importantly, at both time points the specific infectivity of WSN M2 ALPS and WSN M2 RW16 is (statistically significant) reduced compared to WSN M2 Epsin and WSN M2 WT (Fig. 2C).

**Fig. 2:**
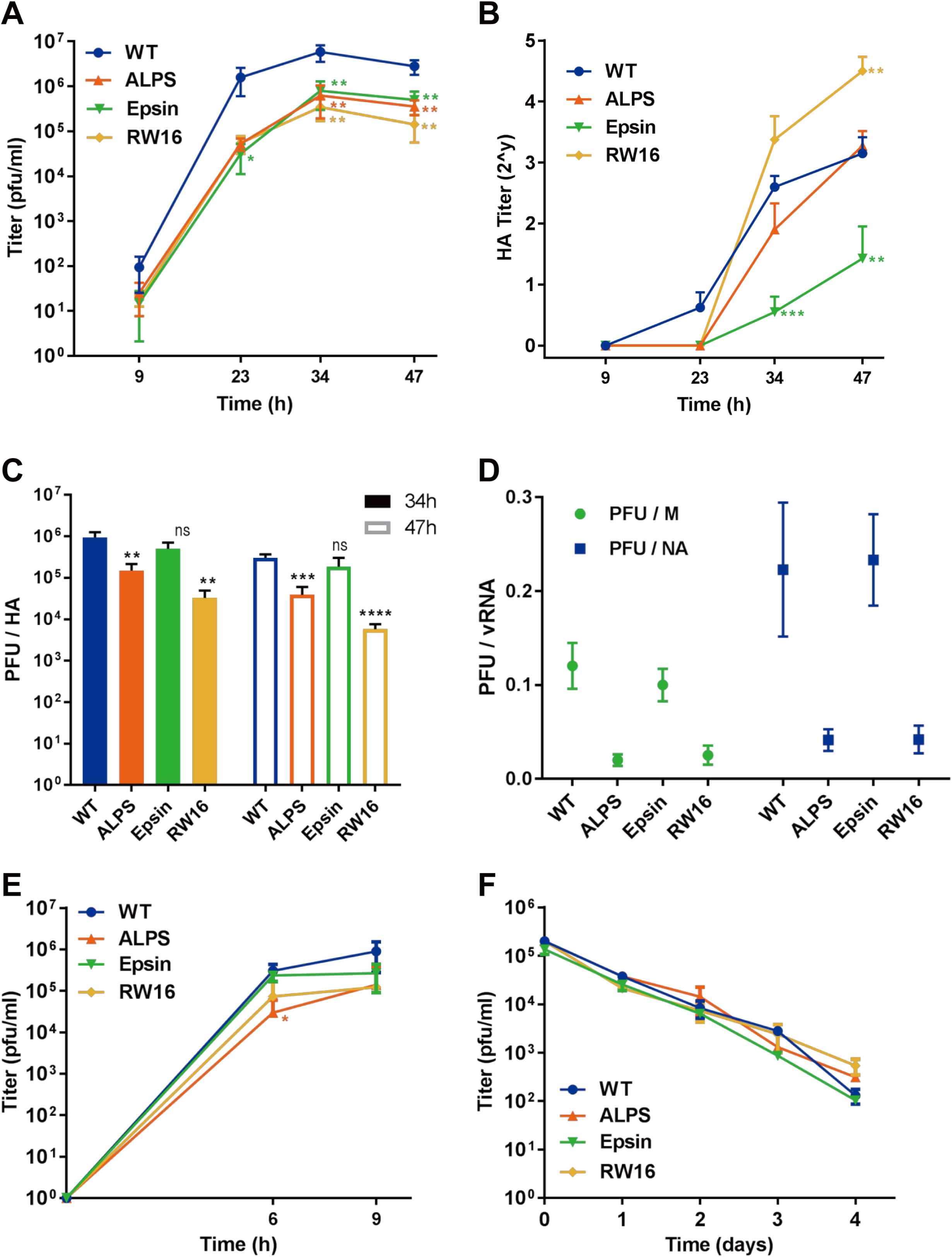
Growth curves, specific infectivity and stability of virus particles. (A-B) Growth curves under multiple cycle growth conditions. MDCK II cells were infected with WSN WT or mutants at a m.o.i of 0.001. Culture supernatants were harvested at the indicated time points after infection and the virus titer was determined by plaque assay (A) or HA-assay (B). Asterisk (*) indicates statistically significant differences between WT and mutants (* P<0.05; ** P<0.01, *** P<0.005) according to a Student’s t-test. (C) Determination of the ratio of infectious to total hemagglutinating particles released at 34h and 47h post infection. Asterisks (*) indicates statistically significant differences between WT and mutants (** P<0.01, *** P<0.005, **** P<0.0005) according to a Student’s t-test. (D) Determination of the ratio of infectious to genome containing particles released at 34h post infection. RNA was extracted from the same volume of culture supernatant. The copy number for gene segment M and NA were determined by RT-qPCR. The ratio of the PFU titers to vRNA copy numbers for three different infections is shown as means±standard deviation for each virus and gene segment. (E) Growth curves under single cycle growth conditions. MDCK II cells were infected with WSN WT or mutants at an m.o.i. of 1 and culture supernatant was harvested at 6h and 9h post infection. Virus titer was determined by plaque assay. The asterisk (*) indicates statistically significant differences between WSN WT and mutant ALPS (* P<0.05). (F) Stability of WSN WT and mutants. 2×10^5^ PFU of the indicated viruses were incubated at 37°C for the indicated time period and its titer was determined by plaque assay.

We then applied RT-qPCR using primers for the gene segments encoding M and NA, respectively to determine the number of total genome-containing particles released at 34h post infection from MDCK cells (Fig. 2D). The determined number was correlated with the infectious virus titer to calculate the ratio of fully infectious to total (genome-containing) particles, which is for wild type virus ∼0.1 if the vRNA encoding M is determined, and ∼0.2 for the NA gene segment. This is at the upper limit for previous estimates for the ratio of total to fully infectious particles which is in the range from 0.1 to 0.01 (52). When compared with WSN WT, WSN M2 Epsin exhibits a very similar ratio, while the other two mutants exhibit a 3-5fold lower proportion of infectious to total particles, regardless of whether the M or NA gene segment were analyzed. In sum, whereas WSN WT and WSN Epsin produce the same ratio of infectious to total particles, the other two mutants produce relatively more non-infectious particles suggesting that the assembly process is less accurate in M2 ALPS and M2 RW16.

In multiple cycle growth experiments a reduced virus titer might be due to a defect in virus budding, in virus entry or in both processes. Therefore, we analyzed virus growth also under one cycle growth conditions by infecting cells with the same m.o.i of 1 and determined virus titers at 6h and 9h post infection. Under those conditions WSN mutants showed no or only slightly (1log) lower titers than WSN WT, statistically significant different from WT only for M2 ALPS at 6h post infection (Fig. 2E). Although defects in virus release might add up after multiple cycles of replication, the result suggests that replacing the amphiphilic helix in M2 causes defects in virus budding; entry of the resulting particles into cells is apparently also affected,.

M2 binds to the autophagy protein LC3 and recruits LC3-conjugated membranes to the viral budding site. This process is required for virus stability, probably because it delivers appropriate lipids to the plasma membrane during budding (14). This is especially evident for filamentous viruses, but the spherical PR8 viruses with a mutation in the LC3-binding domain also show a larger drop in virus titers after incubation at room temperature compared to the corresponding wild type virus. In addition, M2 binds cholesterol (24, 25, 36), a lipid that increases the rigidity of viral membranes and hence probably infectivity of viruses (53–56). To test the possibility that mutant viruses are more unstable than wild type virus we incubated 2×10^6^ infectious virus particles at 37°C for 4 days, removed an aliquot every day and determined the remaining vial infectivity (Fig. 2F). However, no difference was observed between WSN WT and mutant viruses, virus infectivity dropped by ∼1log per day in all cases.

### Little evidence for impaired membrane scission by replacement of M2’s amphiphilic helix

Defects in budding might be reflected by aberrant morphology of the resulting virus particles. Especially if the last budding step (scission) is disturbed this might led to a ‘beads-on-a-string’ morphology, in which individual virions fail to be separated from each other and/or from the plasma membrane (35, 37). To examine whether the cellular helices present in M2 have an influence on virus budding, we applied transmission electron microscopy on ultrathin sections (∼70nm) of MDCK II cells infected at a low m.o.i and prepared 24-36h after infection. Around 60 (WSN Epsin), 120 (WSN RW16) and 280 (WSN Alps) budding events were inspected, but no such morphology was observed for any of the mutants. However, for WSN RW16 we observed two virus particles still attached by a small neck to filopodia (Fig. 3G), but other particles apparently bud normally (Fig. 3H+I). WSN Epsin forms a few bacilliform particles (Fig. 3D). However, most virus particles were spherical for any mutant, yielding no evidence for drastic perturbation of virus morphology by mutations in M2. Thus, we obtained little evidence that the defects in growth of mutant viruses are due to impaired scission of particles from the plasma membrane of infected cells.

**Fig. 3:**
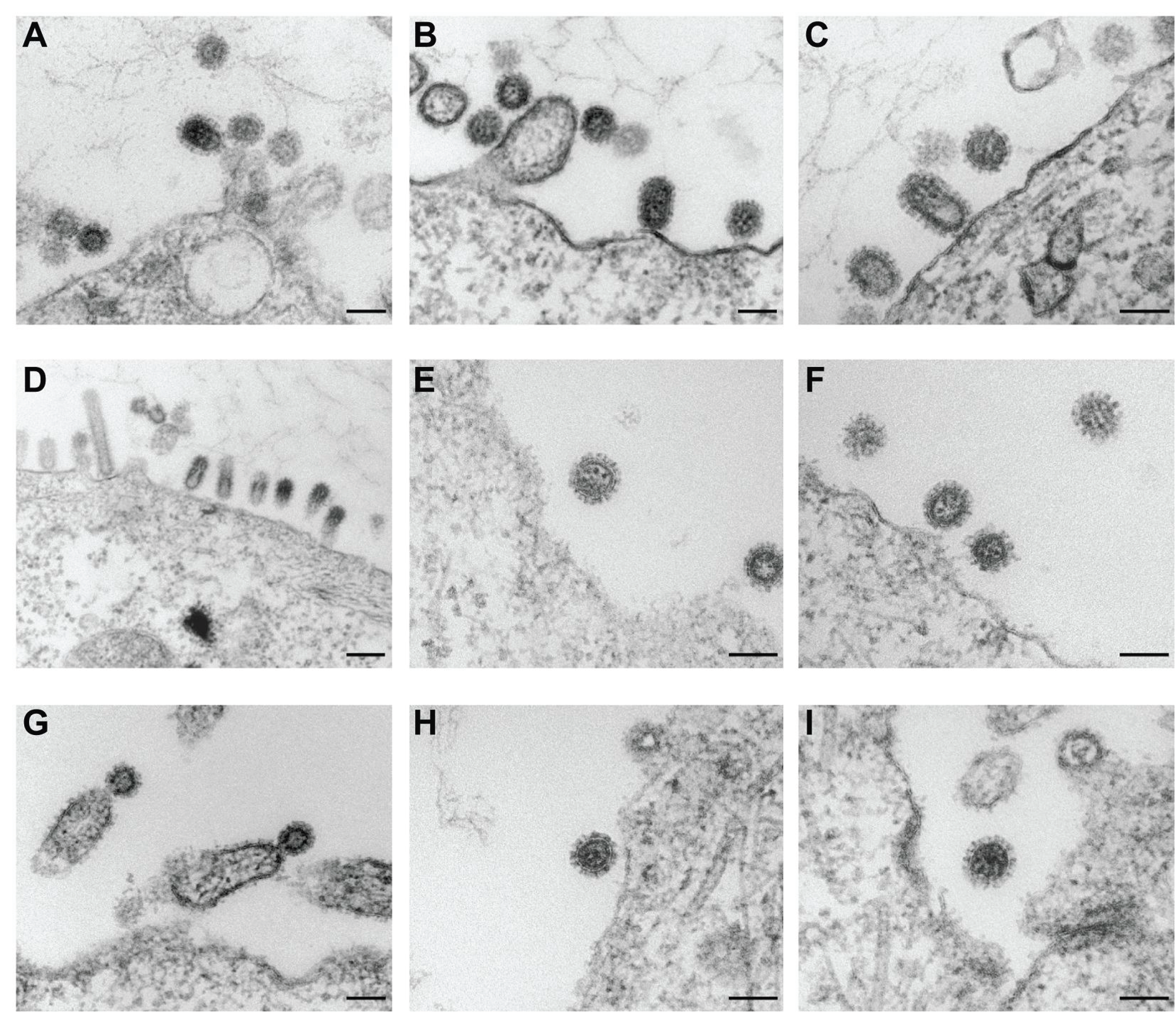
Transmission electron microscopy of infected cells. (A-I): Representative ultrathin sections of MDCK II cells infected with (A) WSN WT, (B-C) WSN ALPS, (D-F) WSN Epsin and (G-I) WSN RW16. Scale bars: 100 nm.

### M2 ALPS and M2 Epsin are less abundantly expressed at the plasma membrane

One reason for the reduced virus titers might be a lower availability of mutant M2 at the budding site. To compare the intracellular distribution of wild type and mutant M2 proteins we used immunofluorescence of permeabilized MDCK II cells 4.5 hours after infection using anti-M2 mAb 14C2. Wild-type M2 and each mutant is present at the plasma membrane, but cells also revealed bright perinuclear (presumably Golgi) and weaker reticular staining (possibly ER) throughout the cell (Fig. 4A).

**Fig. 4:**
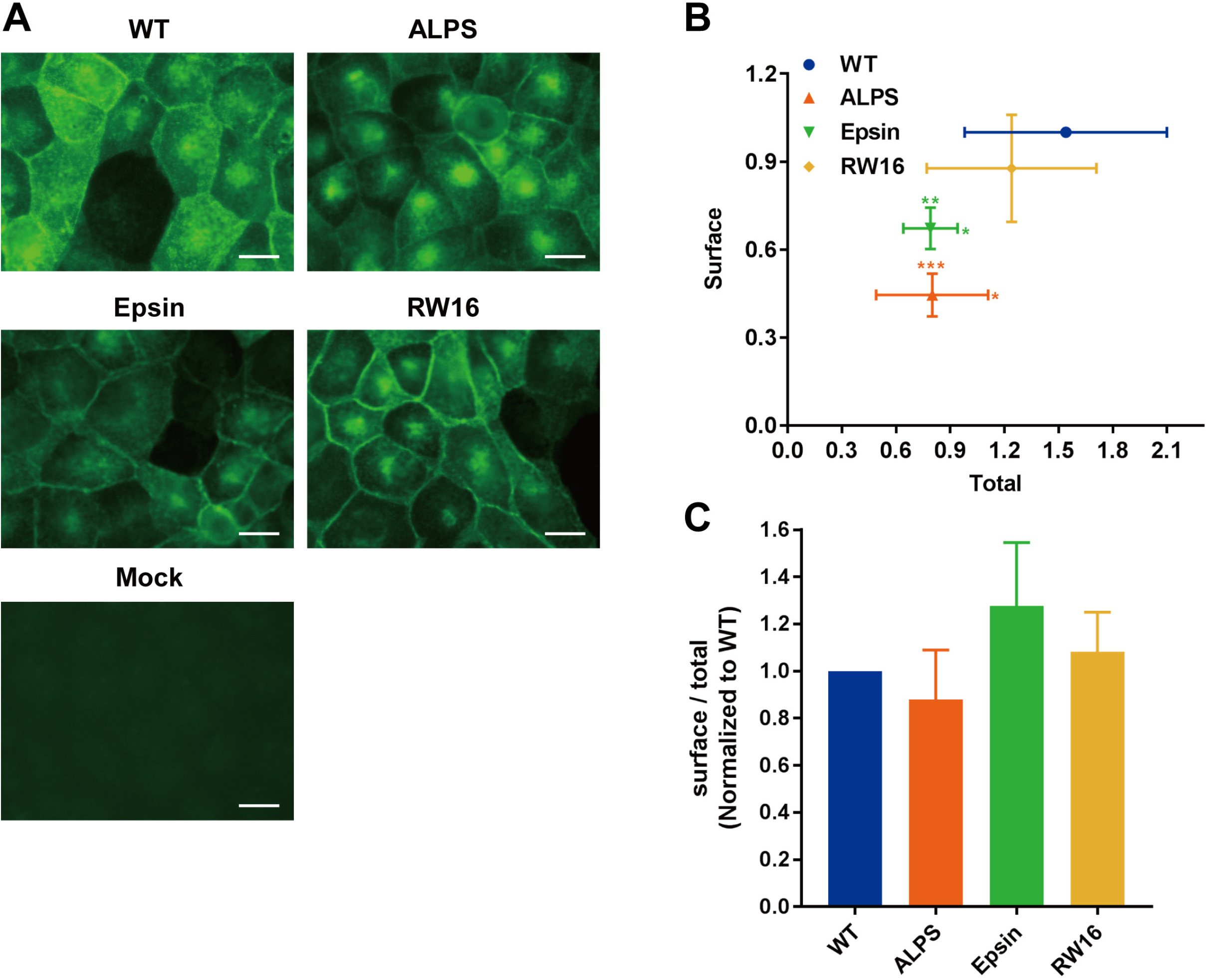
M2 expression and surface transport in infected cells. (A) Fluorescence microcopy of MDCK II cells infected with WSN WT and mutant virus. Cells were infected at a m.o.i of 1, fixed and permeabilized at 4.5h p. i. and stained with anti-M2 mAb14C2 and secondary antibody coupled to Alexa Fluor 488. Note that the images of ALPS and Epsin were generated with longer exposure times. Mock: uninfected cells. Scale bar: 20 µm. (B) Quantification of M2’s expression levels by flow cytometry. Infected MDCK II cells were fixed at 4.5h p. i. and either directly stained with anti-M2 mAb 14C2 (= surface expression) or permeabilized prior to staining (= total expression). The mean fluorescence intensity from at least 100.000 cells was determined by flow cytometry. Results were normalized against surface expression of WSN WT (=1) for each infection and relative surface expression is plotted against total expression for each virus. Results from three individual infections are shown as means ± standard deviation. The asterisks indicate statistically significant differences between WSN WT and mutants ALPS and Epsin (* P<0.05), ** P<0.01, *** P<0.005) according to a Student’s t-test. (C) Calculation of the relative surface expression divided by the total expression from the data shown in (B). A Student’s t-test does not reveal any significant difference between WT and any of the mutants.

We quantified the presence of wild-type and mutant M2 proteins at the cell surface in virus infected cells using antibody staining and flow cytometry. One aliquot of infected MDCK cells was permeabilised to determine total M2 expression levels; the other was left untreated to estimate cell surface fluorescence. Samples were incubated with the M2-antibody 14C2, which recognizes an epitope in the ectodomain of M2 (57), and fluorescent secondary antibody and the mean fluorescence intensity from 10^5^ cells (minus background fluorescence of uninfected cells) was determined. Ratios of total versus surface expression were calculated and results were normalized against the surface expression level of M2-WT (=100%). The resulting graph from four different infection experiments reveal that surface expression of M2 Epsin is (statistically significantly) reduced to 65% and M2 ALPS to 40%. (Fig. 4B). However, this is (manly) due to a reduction in the total expression of both mutants. If the ratio of surface expression to total expression is calculated and normalized to wild type no difference between M2 WT and the M2 mutants is obvious (Fig. 4C).

### Reduced amounts of M2 are incorporated into mutant virus particles

Since especially M2 ALPS and M2 Epsin are less abundantly expressed at the plasma membrane, we asked whether they are less efficiently incorporated into virus particles. We purified WSN WT and WSN mutant virus particles with sucrose gradient centrifugation from MDCK II cells and used western blotting to visualize M2. To quantify the amount of M2 we related the chemiluminescent signal intensity for M2 to that of M1 probed in parallel on the same membrane. The results from three virus preparations (Fig. 5A+B) show that the relative amount of M2 is significantly reduced to ∼25% in M2 ALPS, and M2 Epsin, but also in M2 RW16 relative to M2 WT (=100%).

**Fig. 5:**
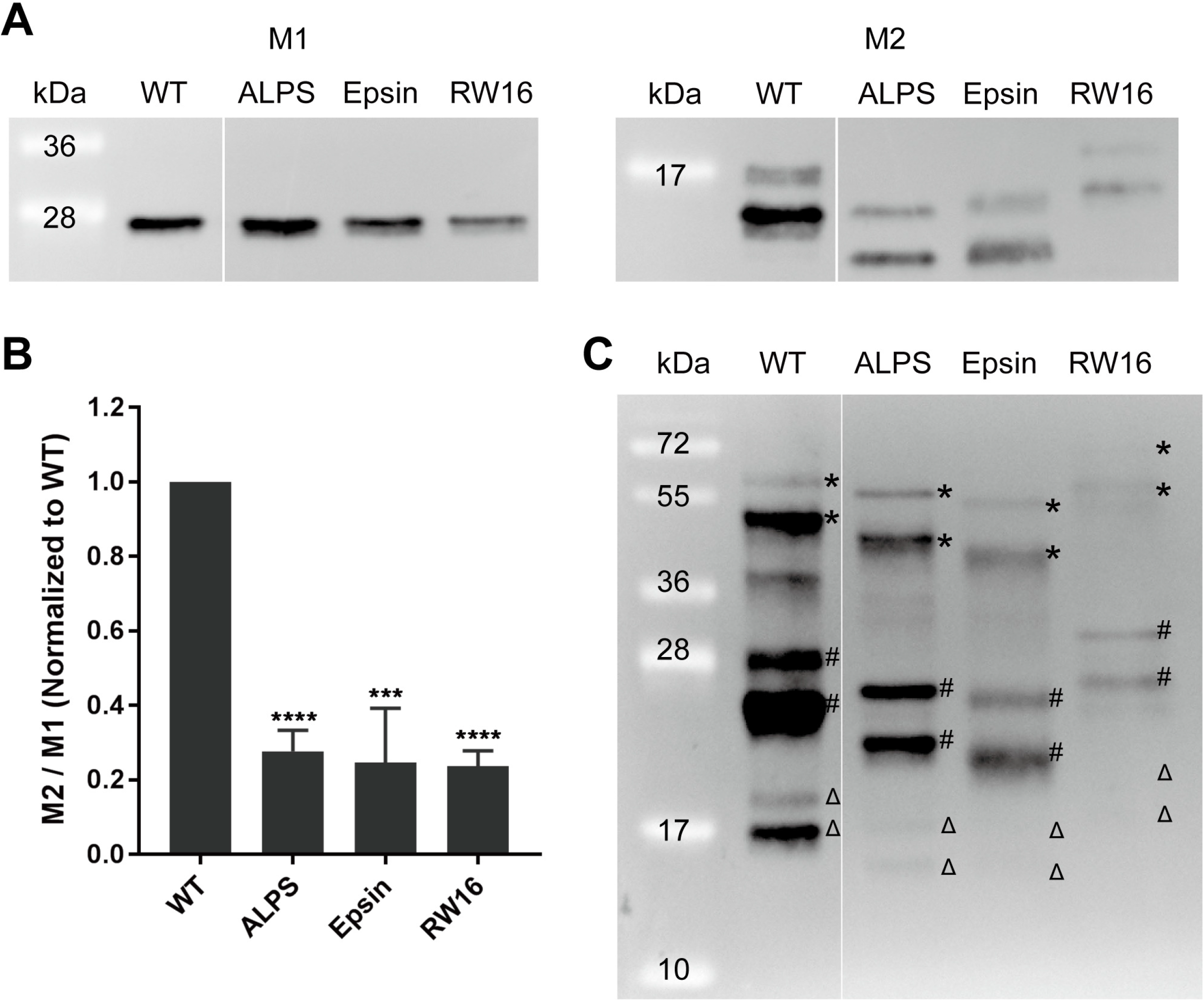
M2 incorporation into virus particles. (A) Gradient purified viruses were subjected to reducing SDS-PAGE and blotted. The membrane was cut between the 28 and 17 kDa molecular mass markers and M1 (left) and M2 (right) were detected by western blotting. The molecular mass markers (kDa) are shown on the left. Samples were equally loaded by volume without any standardization. Determination of the intensity of the two M2 bands revealed that M2 WT has the highest ratio of the lower 15 kDa band relative to the upper 17 kda (4.9, mean of 3 experiments). The ratio is reduced to 2.7 in M2 RW16, to 2.1 in M2 Epsin and to 1.6 in M2 ALPS. (B) Quantification of the ratio of M2 to M1. The density of M1 and M2 bands were determined, the ratio of M2 to M1 were calculated and normalized to WSN WT. Results from three virus preparations are shown as mean±standard deviation. Asterisks indicate statistically significant differences between WT and mutants (*** P<0.005, **** P<0.0005) according to a Student’s t-test. (C) Gradient purified viruses were subjected to non-reducing SDS-PAGE and western-blot with M2 antibodies to analyze oligomerization of M2. Δ: monomer, #: dimer, *: tetramer.

M2 WT appears as two bands under reducing conditions, as observed previously (58). Mutant M2 proteins show a lower ratio of the 15kDa band relative to the 17kDa and a different SDS-PAGE mobility compared to M2 WT. The observed molecular weight does not always correspond to the predicted molecular weight; especially M2 ALPS runs faster than M2 WT although the inserted helix is larger than M2’s authentic helix. Most likely, conformational aspects are involved, since M2 of different virus subtypes having the same number of amino acids run differently in SDS-PAGE (58).

Analyzing M2 proteins by non-reducing SDS-PAGE shows that each mutant forms disulfide-linked dimers and tetramers (Fig. 5C), to a similar extent as M2 WT (5). Thus, except for the slight anomaly in the SDS-PAGE mobility especially of M2 ALPS processing of mutants into tetramers is not disturbed.

### Reduced incorporation of M1 into mutant virus particles

M2 contains a binding site in its cytoplasmic tail to recruit M1 from the Golgi to the plasma membrane (10, 11, 13). To determine whether mutant virus particles contain less M1 we analyzed the protein composition of three purified virus preparations by SDS-PAGE and coomassie staining (Fig. 6A). After non-reducing SDS-PAGE two bands with a molecular weight around 28kDa appeared; both react with M1-specific antibodies in a western-blot (Fig. 6b). Densitometry of the major viral protein bands representing HA, NP and both M1 bands and calculation of their ratios indeed revealed that reduced amounts of M1 were incorporated into each mutant virus particles, most pronounced (and statistically significant) in WSN Epsin and WSN RW16, where it is reduced from 38% (WT) to 16% and 26%, respectively. The reduced amount of M1 is compensated in all mutants by relative higher amounts of HA (Fig. 6C). However, note that assembly of influenza viruses is in general of low fidelity, since even genetically homogenous virus particles released from a single cell show enormous variation in size and protein composition, i. e. the copy number of individual proteins vary up to 100fold between virions (59).

**Fig. 6:**
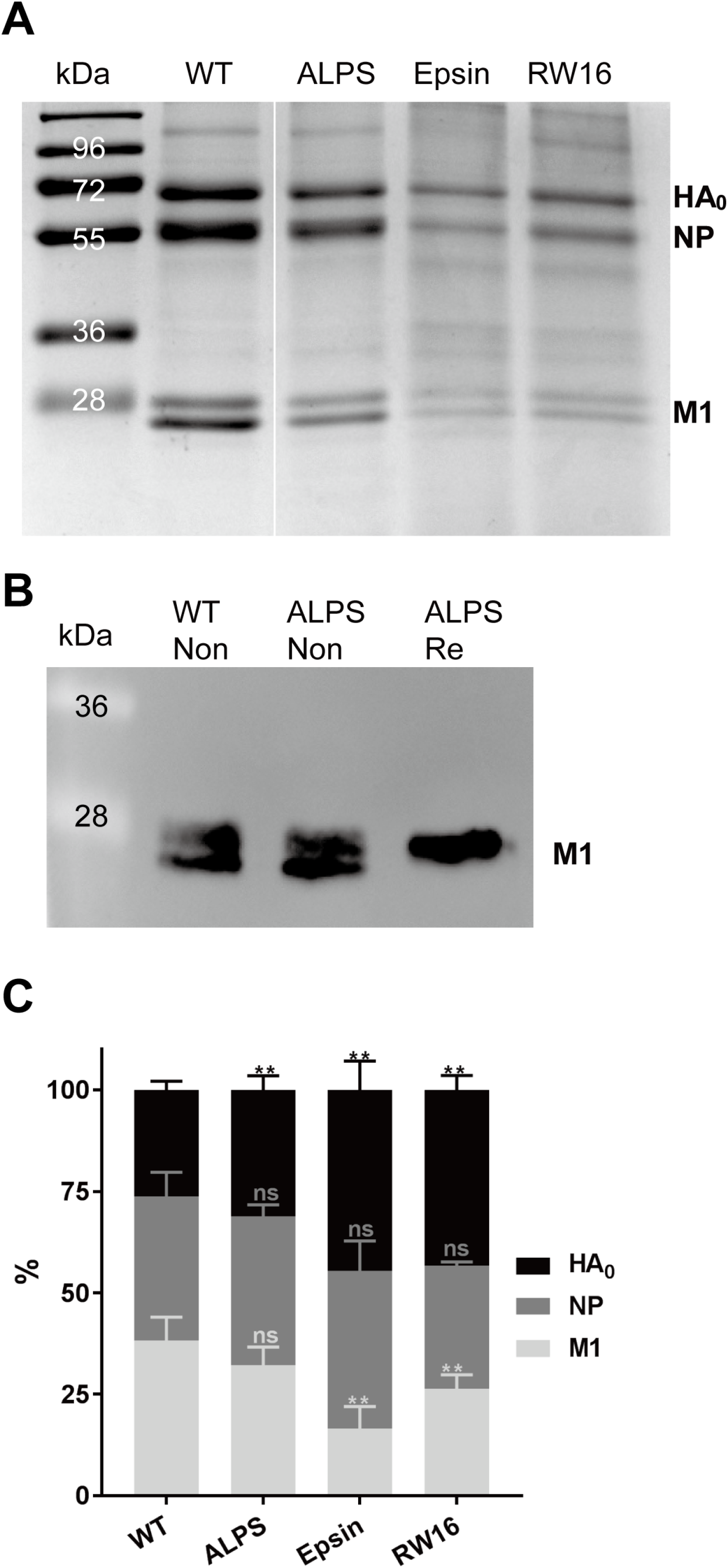
Protein composition of virus particles. (A) Gradient purified viruses were subjected to non-reducing SDS-PAGE and Coomassie staining. The position of HA0, NP and M1 are shown on the right and the molecular mass markers (kDa) are shown on the left. Samples were equally loaded by volume without any standardization. (B) Western-blot of WSN WT and WSN ALPS virus preparations separated under non-reducing (Non) or reducing (Re) conditions using antibodies against M1. (C) Quantification of the relative protein composition. The densities of HA0, NP and both M1 bands from this and two other preparations were determined, and the relative percentage of each protein was calculated. Results are shown as mean ± standard deviation. Two asterisks (** P<0.01) indicates statistically significant differences between WT and mutants according to a Student’s t-test.

### Reduced ion channel activity of M2 ALPS and M2 Epsin

The functional core of M2 consists of the transmembrane region (amino acids 21-51), but the amphiphilic helix is important for ion channel stability and maximal activity (7). To test whether replacement of the amphipathic helix in M2 affects its proton channel activity we used an established assay (60) to measure pH-dependent changes in the fluorescence intensity of the enhanced yellow fluorescent protein (eYFP) in transfected 293T cells which were also infected with Influenza virus. FACS analysis of cells incubated in neutral buffer yielded no change in mean fluorescence intensity (MFI) over time (Fig. 7A). Likewise, transfected, but uninfected cells did not reveal a change after acidification confirming the validity of the assay (Fig. 7B). However, low-pH treatment of infected cells resulted in an obvious decrease in MFI for all Influenza viruses, but the extent of the reduction differed between M2 WT and the mutants WSN Epsin and WSN ALPS. Since the extent of the reduction depends on the number of functional M2 channels on the cell surface, we also determined surface transport by staining of cells with M2 antibodies. All three mutants revealed a small reduction in surface staining (M2 RW16 to 95%, M2 ALPS and M2 Epsin to 80%), which is less pronounced compared to MDCK cells (Fig. 4B). However, the slope of the MFI curve is an intrinsic property of individual proton channels. Whereas WSN WT and WSN RW16 show an exponential decay in MFI in the first three minutes after acidification, it is less distinct in WSN Epsin and the decay is almost linear in M2 ALPS. We conclude, that the proton channel activity of M2 is reduced in WSN Epsin and especially in WSN ALPS, whereas WSN RW16 does not exhibit a defect in ion transport.

**Fig. 7:**
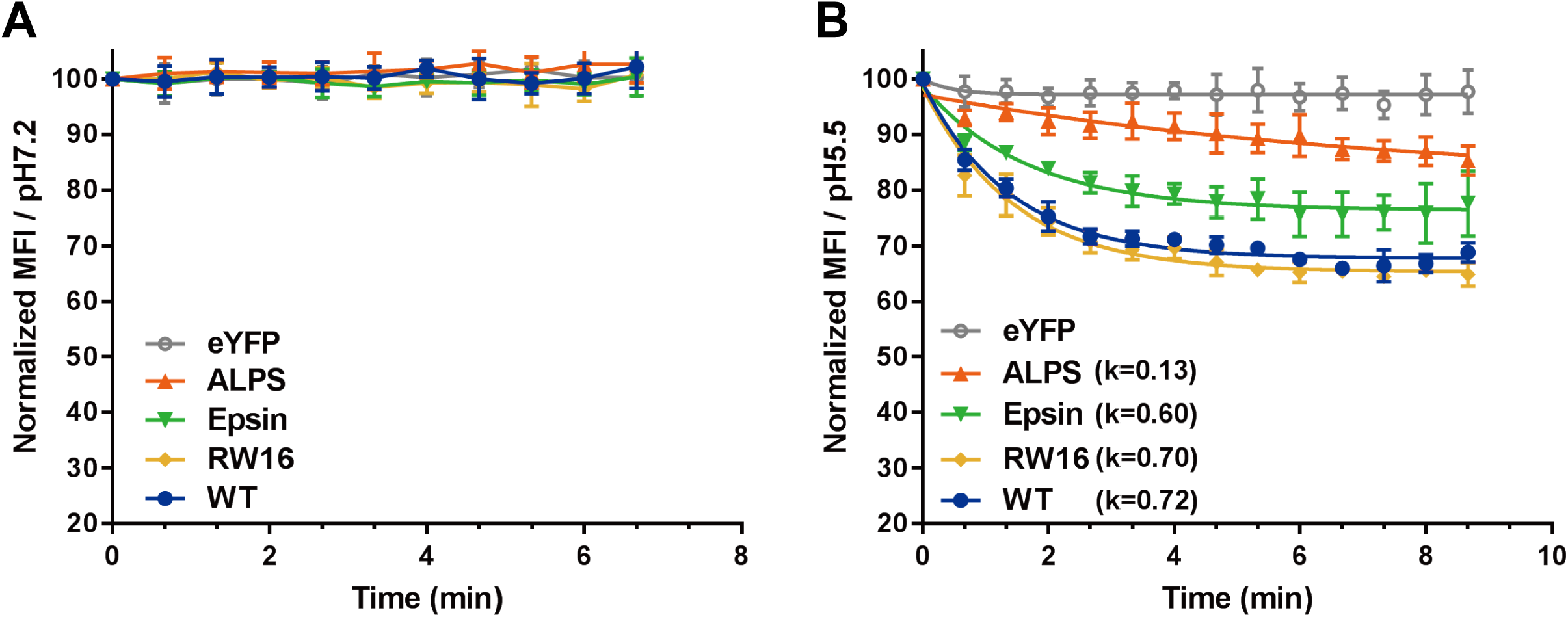
Proton channel activity of M2. 293 cells were transfected with a plasmid encoding eYFP and 24 hours later infected with the indicated influenza viruses. 6 hours later cells were detached from the culture plate, washed and dissolved in either neutral buffer (pH 7.2, A) or acidic buffer (pH 5.5, B) and the fluorescence intensity of at least 10000 cells was measured every 40 second by FACS for 7 (A) or 9 (B) minutes. eYFP: cells transfected with plasmid encoding eYFP. Results from four independent experiments were normalized to WSN WT and the resulting mean fluorescence intensity (MFI) is plotted against the time. Solid lines represent curve fits to one-phase exponential-decay models, the calculated k-values are indicated for each M2. An aliquot of cells was stained with M2 antibodies and calculation of relative surface exposure of M2 revealed a reduction relative to WSN WT (=100%) to 80% for WSN ALPS and WSN Epsin and to 95% for WSN RW16.

## Discussion

In this study, we show that the presence of an amphiphilic helix in the cytoplasmic tail of M2 from the spherical WSN strain is essential for virus replication. Deletion of the helix or replacing it with a scrambled version that has the same amino acid composition but does not exhibit a significant hydrophobic moment, prevented generation of recombinant virus particles. However, the helix could be replaced with three different types of amphiphilic helices with only marginal effects (∼1-2 log reduction) on virus titers. (Fig. 2). Thus, apart from being able to form an amphiphilic helix, there is apparently no strict amino acid sequence requirement in the region proximal to the transmembrane region of M2, as suggested previously (38).

Using ultrathin-section TEM of virus-infected cells we did not observe for any of our mutants virus particles with a bead-on-a-string (and only little evidence for any other altered) morphology as they were observed if bulky residues in the hydrophobic face of the helix of M2 from the Udorn or WSN strain were replaced by alanine (36, 37) or, in general, if viruses with a budding defect were investigated (61) (Fig. 3). This suggests (but does not prove) that virus budding and scission and hence induction of membrane curvature by the mutant M2 proteins is not strongly impaired and thus the reduced virus titers might be also due to other defects in this protein.

Indeed, we observed various other deficiencies in M2 if the helix is replaced, which however differed between the M2 mutants. Two of the mutants, M2 ALPS and M2 Epsin are less abundantly expressed at the cell surface, mainly because their expression levels are decreased (Fig. 4). One might speculate that these mutant M2 proteins are not able to interact with one of the various cellular proteins, which are required to efficiently target M2 to the plasma membrane (36, 62, 63). Assuming that the amount of M2 at the plasma membrane is a limiting factor for the production of infectious virus particles, the reduced surface expression of M2 ALPS and M2 Epsin might partly account for the reduced virus titers. This is consistent with a recent report describing that altering the expression and the intracellular targeting of M2 has major effects on virus replication (64).

Both M2 ALPS and M2 Epsin are also inefficiently incorporated into virus particles, the number of M2 molecules (relative to M1 WT) is reduced to ∼25%. The same reduction was also determined for M2 RW16, although this M2 mutant has no (statistically significant) defect in cell surface transport. One might assume that M2 RW16 is not enriched at the viral budding site and therefore less efficiently incorporated into virus particles. The signals for co-clustering of M2 with HA are not (as initially assumed) palmitoylation and cholesterol-binding to the amphiphilic helix, but other amino acids in its hydrophobic face, such as isoleucine, phenylalanine and tyrosine (37). The helix of M2 RW16 contains only charged (R) and aromatic (W) amino acids and this might prevent privileged access to the viral assembly site. Whether M2 ALPS and M2 Epsin are enriched at the viral budding site cannot be concluded from our data. Note, however, that the α0-helix of Epsin binds to PtdIns(4, 5)P2 even outside of the ETNH-domain and that HA also co-localizes with PtdIns(4, 5)P2 (65). Thus, the proposed mutual affinity of HA and M2 Epsin for this negatively charged lipid present only at the inner leaflet of the plasma membrane might cause their co-localization.

It is not understood why M2 is abundantly expressed in cells, but (in comparison to HA) largely excluded from virus particles (58). One reason for the high expression levels of M2 might be that it interacts with the (also abundantly expressed) M1 protein in the Golgi to transport it by a piggy-back mechanism to the plasma membrane (13). Indeed, we observed that all virus particles having a mutant M2 contain reduced levels of M1, statistically significant for WSN Epsin and WSN RW16, regardless of whether the amount of mutant M2 is also reduced (Fig. 6). Amino acids 71–73, but also residues 45-69 (which encompasses the amphiphilic helix) contain a binding site for M1 (10, 11) and thus replacement of the helix might diminish binding to the matrix protein and hence its transport to the plasma membrane. Furthermore, the ratio of infectious to total virus particles is reduced in the mutants WSN ALPS and WSN RW16 by ∼50%. This result was obtained both by calculating the PFU to HA-titer ratio as well as the PFU to M- and NA-genome segment containing particles (Fig. 2). This suggests that the regular assembly process is somewhat disturbed in the mutants WSN ALPS and WSN RW16 and therefore relatively more non-infectious virus particles are released.

In addition, we determined that the proton channel activity of M2 Epsin and M2 ALPS is also compromised (Fig. 7). This, together with the observation that all mutant virus particles contain a reduced number of M2 proteins (Fig. 5), suggests that they might exhibit defects in virus entry. M2 mediated acidification of the virus interior must occur before the low pH in the endosome triggers HA-mediated membrane fusion. Otherwise the diffusion of protons from the endosome through the fusion pore into the cytosol would be faster than M2-mediated transport into virus particles (66, 67).

In summary, we obtained only little evidence that scission of virus particles is disturbed if the amphiphilic helix of M2 is replaced by helices from cellular proteins having similar biophysical properties. Instead, we observed various other functional deficiencies in individual mutant M2 proteins, such as reduced exposure at the plasma membrane, reduced incorporation into virus particles and assembly defects, which might be responsible for the moderately reduced viral titers. Since we could not generate infectious virus if M2’s helix was deleted or if it was replaced by a scrambled version that does not exhibit a hydrophobic moment, our data support the concept that the amphiphilic helix in M2 inserts into the lipid bilayer to sense membrane curvature at the neck of budding viruses and/or to induce membrane curvature that causes scission of virus particles. This is consistent with the concept that M2 is member of a family of membrane scission proteins, which have similar biophysical properties, but no homology in the amino acid sequence (31).

## Acknowledgements

This work was supported by the Human Frontiers Science Program (HFSP). Bodan Hu is recipient of a PhD fellowship from the China Scholarship Council (CSC). The funders had no role in study design, data collection and interpretation, or the decision to submit the work for publication.

